# Generative adversarial neural networks maintain decoder accuracy during signal disruption in simulated long-term recordings from brain computer interfaces

**DOI:** 10.1101/2021.05.18.444641

**Authors:** Thomas Stephens, Jon Cafaro, Ryan MacRae, Stephen Simons

## Abstract

Chronically implanted brain-computer interfaces (BCIs) provide amazing opportunities to those living with disability and for the treatment of chronic disorders of the nervous system. However, this potential has yet to be fully realized in part due to the lack of stability in measured signals over time. Signal disruption stems from multiple sources including mechanical failure of the interface, changes in neuron health, and glial encapsulation of the electrodes that alter the impedance. In this study we present an algorithmic solution to the problem of long-term signal disruption in chronically implanted neural interfaces. Our approach utilizes a generative adversarial network (GAN), based on the original Unsupervised Image to Image Translation (UNIT) algorithm, which learns how to recover degraded signals back to their analogous non-disrupted (“clean”) exemplars measured at the time of implant. We demonstrate that this approach can reliably recover simulated signals in two types of commonly used neural interfaces: multi-electrode arrays (MEA), and electrocorticography (ECoG). To test the accuracy of signal recovery we employ a common BCI paradigm wherein a classification algorithm (neural decoder) is trained on the starting (non-disrupted) set of signals. Performance of the decoder demonstrates expected failure over time as the signal disruption accumulates. In simulated MEA experiments, our approach recovers decoder accuracy to >90% when as many as 13/ 32 channels are lost, or as many as 28/32 channels have their neural responses altered. In simulated ECoG experiments, our approach shows stabilization of the neural decoder indefinitely with decoder accuracies >95% over simulated lifetimes of over 1 year. Our results suggest that these types of neural networks can provide a useful tool to improve the long-term utility of chronically implanted neural interfaces.

## Introduction

Implantable neural interfaces have been used experimentally in animal models and medically in humans to support brain-computer interfaces (BCIs). These interfaces record neural activity and then typically try to decode its meaning to control external devices. For example, neural activity measured in the motor cortex of a man with tetraplegia has been used to guide a computer cursor and robotic arm. More broadly, many diverse goals can be served by developing the ability to measure and accurately decode neural activity [1]. Neural decoder training requires measuring responses to large-scale, labeled training sets, making it a laborious and costly process [2, 3]. To avoid repeating this process, decoders would ideally be trained using neural activity measured shortly after the neural interface is implanted and then, have their trained parameters fixed for long-term decoding. However, long-term decoder functionality is undermined by changes in the neural activity measured by the BCI [4]. The failure of BCI decoders to function long-term significantly limits their utility in real-world applications. In this paper we describe an algorithmic approach to correcting these changes in the observed neural activity in simulated chronic recordings across two different types of neural interfaces.

Changes in the neural activity measured on a BCI are caused both by biological and material changes and will depend on the type of BCI used. Two common implantable devices used for BCIs are multielectrode arrays (MEAs) and Electrocorticography (ECoG) arrays. MEAs are often implanted on the cortical surface of the brain and can measure action potentials (spikes) from local populations of neurons. Spikes measured on each electrode, or assigned to individual neurons, are often decoded by their event frequency in peristimulus time histograms (PSTHs). The observed spikes measured with MEAs can change for many reasons including: electrode malfunction or movement, local neuron death or adaptation [5-18]. These changes can lead to disruption of the observed stimulus responses that include electrode channels that are lost entirely or more subtle differences in the stimulus-specific spike rates. In contrast to MEAs, ECoG arrays do not measure spikes from individual neurons but instead measure correlated neural activity from large populations of neurons in local field potentials (LFPs). LFPs measured from ECoG can change their power spectrum over time, caused by changes in electrode impedance [19, 20] and cortical adaptation [21, 22]. Changes in the measured LFPs and PSTHs often preserve decodable information but cannot be decoded by the previously trained decoder because the information has moved to a new signal domain. To maintain the decodability of signals chronically in the neural interface, we treat the problem as a domain translation problem. The objective of our approach is the unsupervised remapping or translation of “disrupted” signals to their original “clean” domain before application of an existing trained decoder.

Remapping across data domains has been successfully accomplished using generative adversarial networks (GANs) [23]. GANs are a class of machine learning (ML) networks that can generate data with similar statistics to their training set. Previous work demonstrated use of a GAN to perform unsupervised image-to-image translation (UNIT) and successfully mapped image data from one domain to another (e.g. satellite to street-map images)[24]. The UNIT architecture uses a pair of autoencoders to learn the statistics of the individual data domains, and a shared latent space to facilitate cross-domain translation. Conceptualizing clean and disrupted BCI data as distinct dataset domains, we modified a UNIT architecture to remap disrupted BCI data back to the clean data domain in order to prevent decoding errors. To preserve the decodable information, the network must remap the disrupted data while maintaining its true stimulus identity (i.e. “class”). To reliably maintain class identity, we apply new objective training functions and post-training network selection criteria to the UNIT architecture. We call the networks developed to remap the BCI data, Adversarial Networks for Neural Interfaces (ANNI).

ANNI was trained and tested using simulated data recordings from MEAs and ECoG arrays before and after disruption. A neural decoder was fit to each clean dataset and then tested after disruption, either with or without prior remapping using ANNI. In the absence of ANNI, the neural decoders exhibit large drops in decoder performance with even relatively small data disruptions which are characteristic of failure modes in chronically implanted neural interfaces. However, following remapping with ANNI, decoder performance is significantly increased, often attaining accuracy equal to the clean data. Our results suggest ANNI’s potential to maintain decoder performance and extend the functionality of BCIs for long-term use.

## Methods

### MEA simulation

We use a simple model that mimics the key components of neural responses recorded from an MEA (Figure 1). The model simulates a set of neurons with diverse stimulus responses. First, the response of a single neuron is determined by its response probability to a given stimulus. Response probabilities were determined by heterogenous gaussian tuning curves. The tuning curves’ mean and standard deviations (SDs) were randomly selected within the stimulus range (for mean) or half the stimulus range (for SD). The tuning curves generate response probabilities for each neuron from stimuli values defined in time (400 ms with 1 ms sample rate).

**Figure 1.**
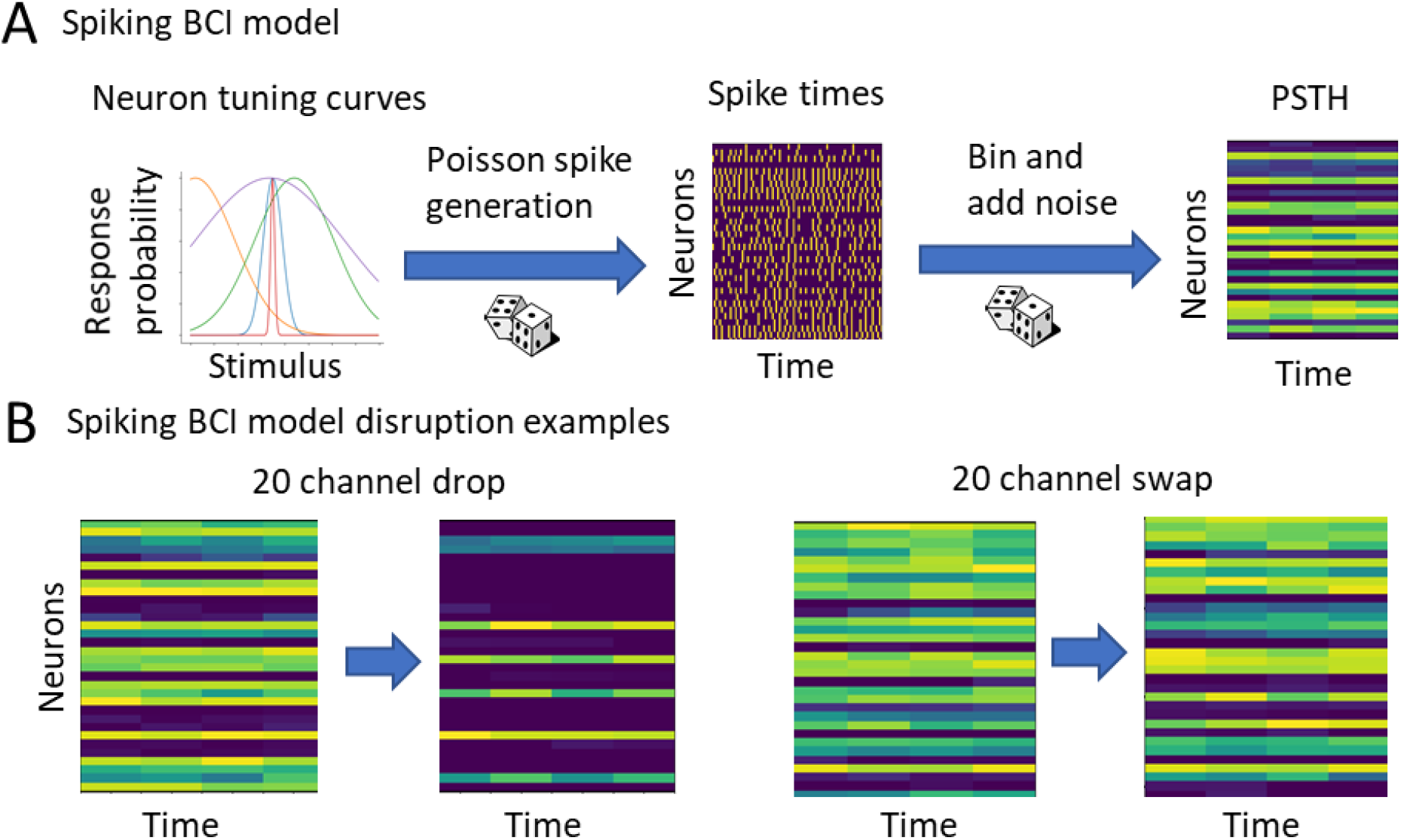
A neural model was used to simulate PSTH datasets that were disrupted via channel drop or swap. **(A)** 32 different tuning curves (6 shown) describe the response probability for a particular stimulus value. Stimuli are static in time at 1 of 20 stimulus values. Variability is injected both in the Poisson generation of spike times and added directly to the PSTH. Thus, PSTHs variably encode 1 of 20 different stimulus values. **(B)** Two different models were used to simulate neural disruption. In an example of disruption, PSTHs have either 20 channels dropped or 20 channels swapped. Noise on all PSTHs, including disrupted PSTHs, is independent.

The stimuli in these experiments were static in time at 1 of 20 different values evenly distributed across the stimulus range. This process generates temporally constant response probabilities. Second, response probabilities were used to generate spikes using a Poisson process to simulate noise in spike generation [2]. Third, spikes counts were binned (100 ms bins), generating 32×4 bin PSTHs (# channels x # time bins). Finally, a second additive noise source was added to the PSTHs. The additive noise had a SD of 10% of the PSTH mean and all resulting PSTH values below zero were rectified at zero. This process generated 5000 PSTH trials encoding 20 different stimulus classes.

### MEA disruption simulation

Recovery was tested from channel drop-out (drop) or neuron drop-in (swap) disruption models. In the drop-out model of neural disruption, each degradation step was defined by the loss of an entire electrode channel, which is simulated by zeroing out all time bins in the PSTH for that particular channel in all samples. This loss is cumulative across the multiple degradation steps such that at step 16, half of the channels contain only zeros. In contrast, a degradation step in the drop-in model was defined by a change in the stimulus response at an existing channel in the set. The stimulus response change was modeled as a 10% positive or negative shift in the tuning curve mean (relative to the entire stimulus range) and an increase or decrease in the tuning curve SD of 2x the original SD. Both models mimic common types of disruptions in MEA recordings and are illustrated by examples in Figure 1B.

### ECoG simulation

We model ECoG at the level of the LFP signal observed at the electrodes rather than the underlying activity of neurons. The reason for this is that forward models of observed activity at the ECoG sensor derived from the underlying neuron populations will be specific to the neural subsystem being measured (e.g. somatosensory vs visual cortex) as well as the type and density of neural array. By simulating a more generic LFP signal and noise, we sidestep the implementation of more complex physical models required for simulating neuronal activity and point spread functions to approximate specific population responses (i.e.[25, 26]) while still replicating the fundamental signal qualities of ECoG reported in the literature (e.g. the frequency spectra, amplitude and noise characteristics)[20, 22, 27, 28].

To model ECoG, we assume that the basic time-domain response at any electrode (channel) will have an S = 1/F distribution in the time-frequency domain, where S is the signal at a given channel and F is frequency in hertz. This spectral distribution is well characterized for ECoG [27, 28]. We further assume that because ECoG samples from a large cortical territory that responses will have spatial heterogeneity. Based on examples in the literature we also assume that changes in the observed ECoG signal that are relevant for decoding typically manifest themselves as changes in lower frequency oscillations when averaged over multiple samples [20].

We simulated LFP signals on an 8×8 mECoG array. A given stimulus (class) is instantiated with a series of signal inputs. Each signal input requires selecting a channel, frequency, phase, time (within the 1s signal window), and amplitude of sinusoidal input. The final parameter controls the spatial distance between electrodes in the array. In practice we simulate this with a unit grid (1mm spacing) and scale the modulating input strength as a function of the distance from the center locus of the modulating input. Each modulating input is added to the 1/F noise at the appropriate channels to produce a spatiotemporal pattern of activity that serves as the starting stimulus class template. Any of these parameters can be manipulated to investigate a different aspect of the model (i.e. array size, frequency of signal inputs etc.). In practice, we have used the following parameters to produce a reasonable sample set: # of signal inputs per stimulus class = 5, frequency range = 3:12 Hz, phase = randomly assigned, amplitude = 1/F, time = randomly assigned. An example of the appearance of these simulated data is shown in Figure 2.

**Figure 2.**
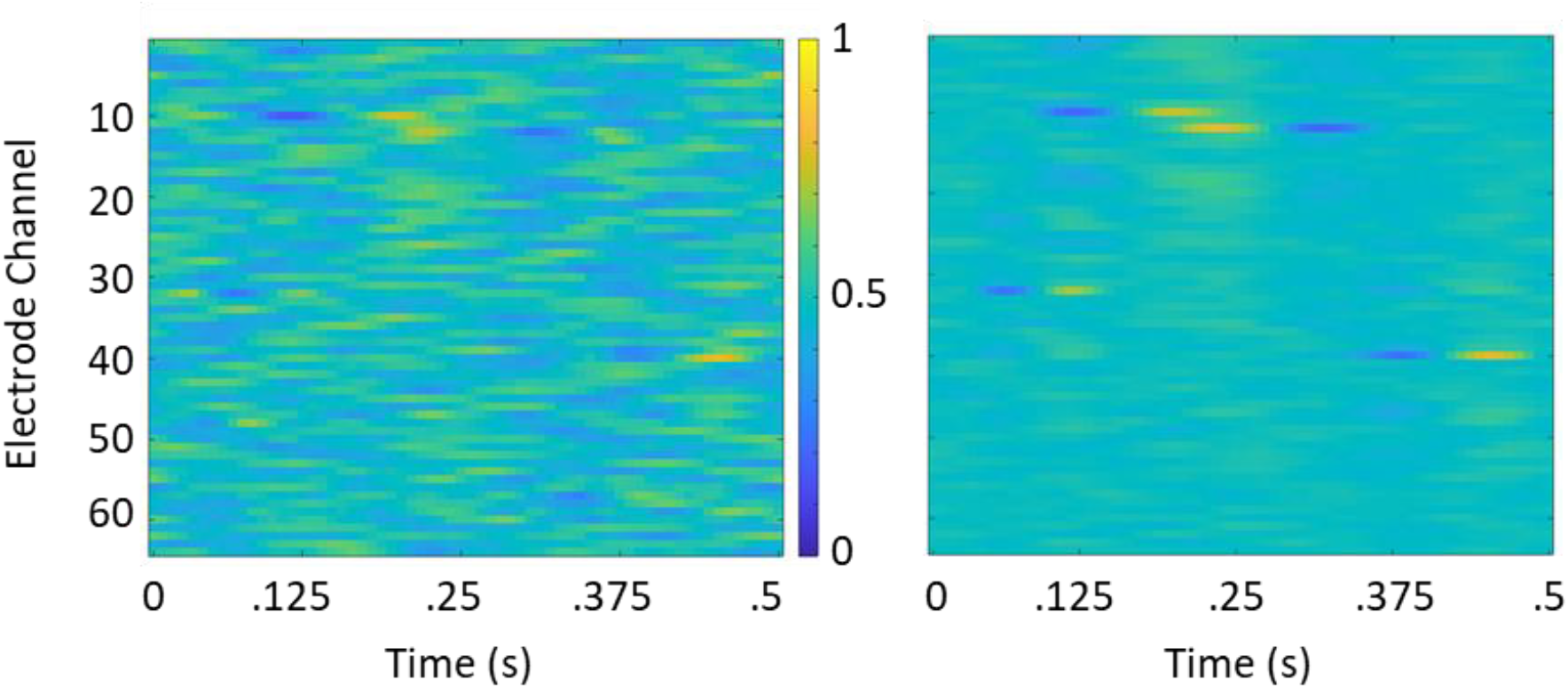
Sample simulated ECoG signals show typical time frequency signal and noise characteristics. **(A)** A single sample of 64 channels and 64 time bins (128 Hz sampling). Signals have been normalized across the entire set from 0:1 for use in neural network training. Signals are barely visible embedded in 1/F noise patterns on each channel. **(B)** Grand average of 25 samples produces easily recognizable signals on channels 10, 12, 32, and 40 which make up the identity for this signal class.

### ECoG disruption simulation

As described above, the primary source of disruption on ECoG is changes in electrode impedance. These changes are particularly egregious during the first 75 days after implant and lead to dramatic changes in the variance and resulting spectral power of signals. We use the linear model fits established by Ung et al. to establish the degree to which changes manifest in each spectral band of the signal over the first 100 days [21]. These changes range anywhere from -0.013 to -0.003 per day depending on the spectral band and the individual from the study.

To simulate changes in spectral power over time we simply need to know the length of time since implant. Changes in expected signals are then computed by applying a discrete Fourier transform to each time-domain signal (e.g for each electrode channel and sample), applying a transform function to the signals in Fourier space with the appropriate frequency band coefficients [21], and then using an inverse Fourier transform to restore the signals to the time domain. Beyond the first 100 days after implant, the interface stabilizes somewhat and the day-to-day or week-to-week changes in signal strength are confined to a narrower operating range [21]. In our simulations we model this operating range as +/-20% of the stable signal strength beyond day 100. These fluctuations are generated randomly and drawn from a gaussian distribution centered at 0% signal change at each time step within the allowable bounds.

The second form of simulated disruption is cortical adaptation. Adaptation refers to changes in the neural responses themselves as the brain learns to more efficiently utilize the interface. The use of ECoG in chronic recordings and brain computer interfaces (BCIs) is limited currently and has restricted the study of adaptation to only a few studies. Those studies suggest that cortical adaptation manifests primarily as an increase in the amplitude of these signals over time [29]. We allow the signals to adapt continuously over the simulated lifetime of the implant in the following way. At each time point in the simulation (10 day steps), we allow 20% of the modulatory input signals to adapt, selected at random. Each time a signal adapts it can increase or decrease its amplitude by 10-20% prior to accounting for the impedance change. The reason for including a long-term depression (signal decrease) mechanism in our simulations was to prevent signal saturation over the longer simulation times (e.g. 360 days). We adapt signals at each time point (every 10 days for a total of 36) in the simulation and signals can be adapted multiple times over the time steps. The absolute magnitude of possible decrease or increase in a signal is capped at 70 – 150% of its starting amplitude over the lifetime of the simulation. The result is a set of signals that continuously change their absolute and relative magnitude to each other over the lifetime of the simulation.

### ANNI System Overview

Figure 3 illustrates the concept of operation for ANNI. Clean signals corresponding to many distinct neural activity patterns are simulated and used to train a neural decoder. Subsequent signals corresponding to the same activity patterns become disrupted over time, as discussed above. For each disrupted dataset the ANNI system trains multiple recovery models, each attempting to recover the disrupted dataset back to its clean state. The reason to train multiple^1^ recovery models on the same dataset is that we have observed high variance in the performance of recovery across random weight initializations and hyperparameters. For each dataset, the model exhibiting the highest decoder entropy is selected as the best model and is immediately applied to incoming neural signals.

**Figure 3.**
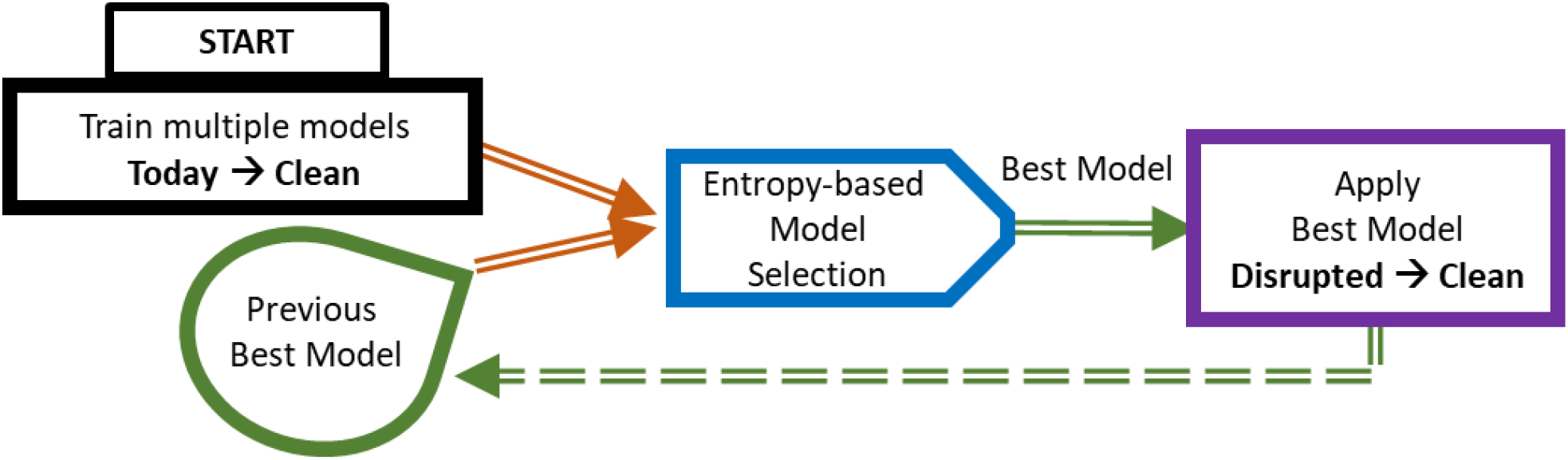
ANNI System Overview. Today’s collected neural recordings are used to train multiple recovery models each attempting to do the same thing – return today’s signals to their undisrupted counterparts. Multiple models are trained due to the high variance in remapping performance. An entropy-based model selection criterion identifies the best among multiple models, and the best model is then applied to incoming neural recordings. As new neural recordings are collected, the former best model (yesterday’s best model) is allowed to compete with newly-trained models. All results reported in Sections 4 & 5 are discussed in the context of this competitive model continuation paradigm.

### Entropy-based Model Selection

As there are no class labels on incoming, disrupted signals, it is impossible to directly assess the recovery of any given signal. We observed that low performing models often lead to decoding fewer class types (e.g. through mode collapse), generating less uniform predicted class distributions. The uniformity may be quantified by the decoded class entropy (see Figure 4). Given a set of possible classes is C, the decoded class entropy H is computed as a sum over each class c ∈ C

**Figure 4.**
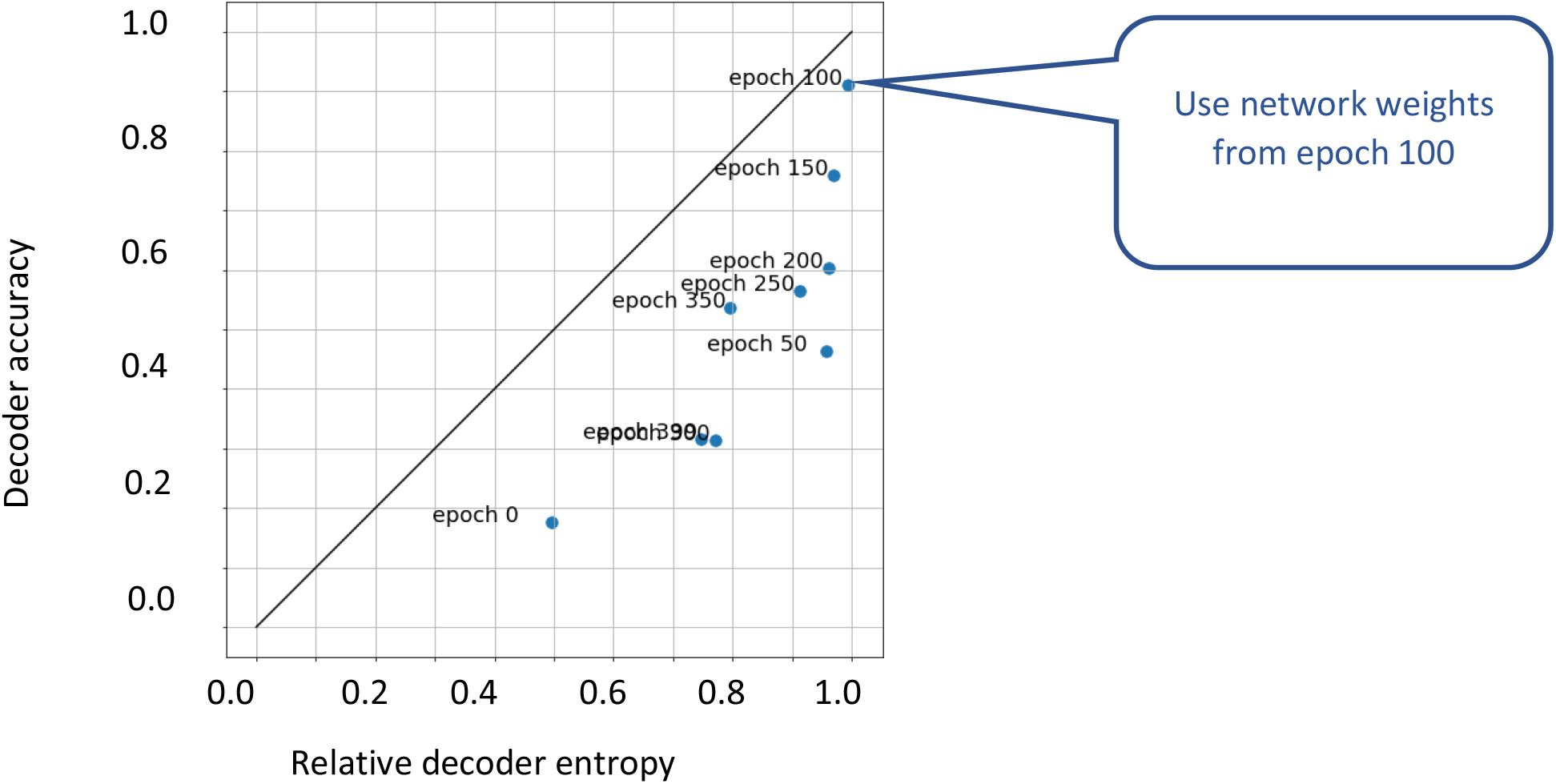
High decoder entropy is well correlated with high decoder accuracy. Each point on this plot is (normalized) decoder entropy vs decoder accuracy at a single epoch during the training of a single recovery model. Because our paradigm assumes no ground-truth class labels during training, we must use a surrogate metric to evaluate recovery performance. In order to identify the network state that yields the highest accuracy model, we select the network weights from the epoch corresponding to the highest decoder entropy.

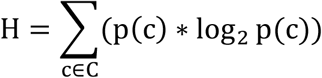

where p(c) is the area-normalized histogram of the decoder class predictions. This criterion assumes that the set of sampled disrupted data contains a representative sampling of the various classes. However, in practice, other selection criteria such as matching recovered signal variances (to clean signals) has worked nearly as well. It is important to note this metric does not need ground truth on the class prediction.

The strong correlation between high decoder entropy and high decoder accuracy at the population level enables the selection of top performers from among multiple models trained to recover the same set back to clean. Figure 4 shows the relationship between decoder entropy and decoder accuracy at the population level – where the population spans the intermediate states of network weights during the training of a single model. The correlation across models each starting from different initial weight initializations and hyperparameters enjoys the same shape as the trend in Figure 4. Thus, our entropy-based selection criteria is used not only to select the best stopping point for a given model during its training, but also to select the best overall model from any number of trained models at a given disruption level.

### Model Continuation

The model selected to recover the current disrupted dataset may also succeed at recovering subsequent disrupted datasets. Thus, we re-submit the previous recovery model for entropy selection with the batch of models trained on the current disruption dataset. In this way, a single previously-trained model could outperform all current models and continue onward indefinitely (we occasionally see single models win out over several disruption steps in our experiments). Our results in Sections 4 & 5 include recovery using the model continuation paradigm.

### Signal Recovery as Domain Adaptation

The key insight of our machine learning signal recovery approach is recognizing that having access to clean neural recordings sets up a domain adaptation problem from disrupted recordings back to their clean state. The key challenge of signal recovery (or domain adaptation) in our setting is to be able to recover individual disrupted neural recordings from an unknown class to their non-disrupted counterparts that are decodable into the correct class by the neural decoder. We base our approach on the UNIT architecture [24] but alter the training objectives to achieve unsupervised and class-dependent domain adaptation needed in the present context. This results in an algorithm that differs from UNIT and other current multi-modal unsupervised domain adaptation approaches [30, 31] in that the resulting class of the transformed signal matches the incoming signal’s class, without having knowledge of the identity of the incoming class beforehand. It is important to note that while any level of disruption affects the same channels of the interface (across all class signals), it is also the case that the information in each signal class will be disrupted in a different way depending on whether that channel carries information that is relevant to its class identity. That is, the same channel disruption affects each signal class differently. This means that ANNI cannot perform a single domain transfer, but instead must perform a class-dependent transformation. For example, it is not enough for the algorithm to learn to transform photographs of both buildings and landscapes into the style of Monet – rather ANNI must learn to transfer buildings into Monet and landscapes into Seurat. This represents a more challenging problem than what has been demonstrated with current unsupervised domain transfer algorithms.

### Architecture Rationale

We adopt the system architecture presented in UNIT because it is a composition of several well-understood neural network modules designed to enable simple, flexible, and intuitive information flow during training and subsequent application. The architecture is illustrated in Figure 5 and consists of two encoder/decoder pairs, one pair forming an autoencoder for clean data and the other pair forming an autoencoder for disrupted data. Both encoders project their incoming signals into a shared latent space, presenting the opportunity to encourage the alignment of the latent code of an undisrupted signal and the latent code of its disrupted counterpart. Additionally, the architecture includes two discriminators (from the GAN paradigm) that present the opportunity to recover disrupted signals without a non-disrupted counterpart. This is essential because any non-disrupted portions of subsequently recorded signals will not match any previously collected signal due to natural variability in neural responses. The generator portion of the GAN analogy is realized by passing a disrupted signal through the encoder Enc_B_ and reconstructing the signal from the latent code z_B_ using Dec_A_. The result x_BA_ ≔ Dec_A_(Enc_B_(x_B_)) must match the statistical properties of the non-disrupted signals. The non-disrupted discriminator will not enforce the class alignment in the non-disrupted signal space – that ability comes from two cycle consistency constraints (discussed below). The architecture is symmetric about the datasets in the sense that passing a signal x_B_ from dataset X_B_ has a counterpart action that begins with passing a signal x_A_ from the dataset X_A_.

**Figure 5.**
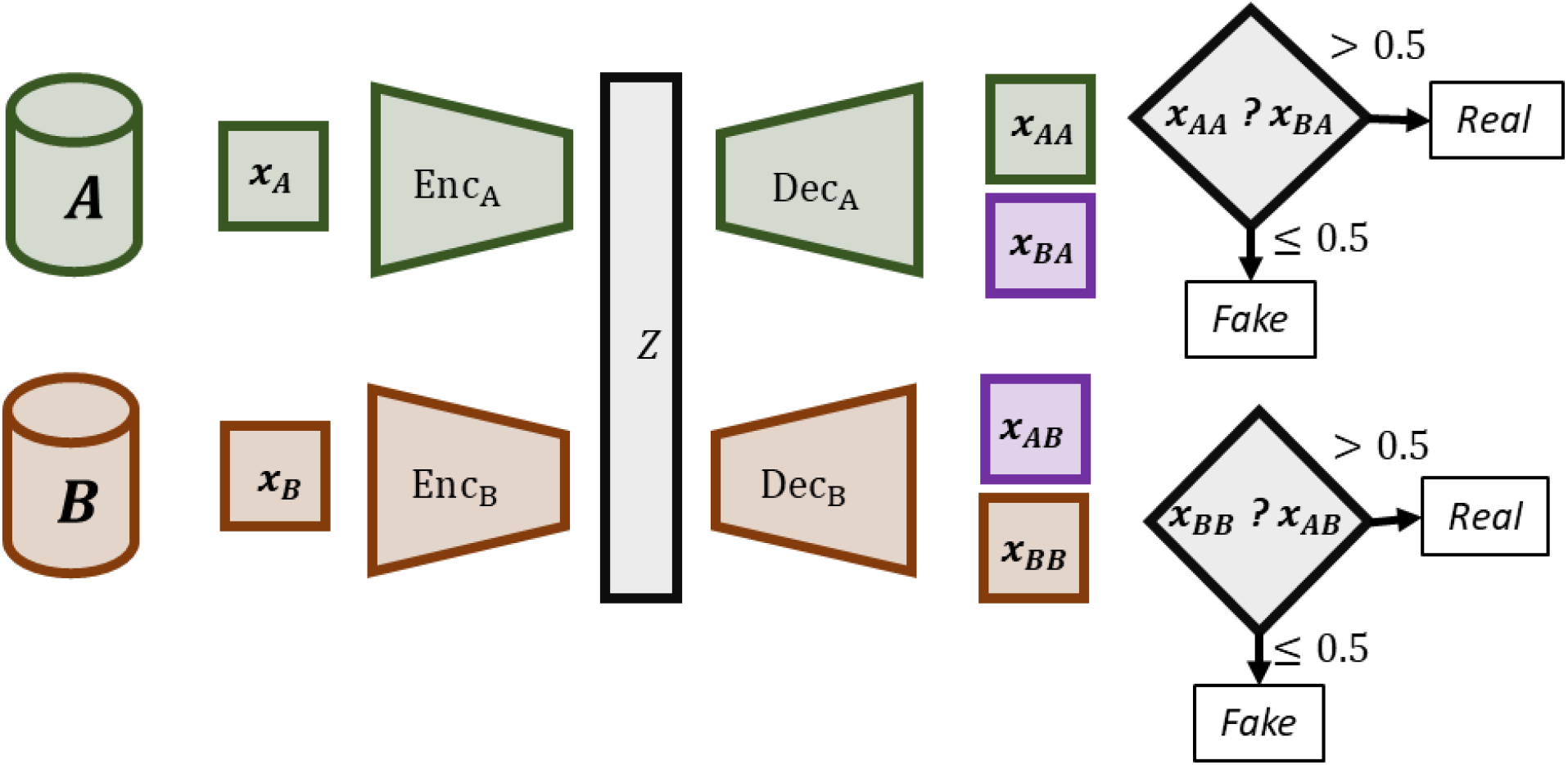
ANNI Neural Network Architecture. Dataset ***A*** contains clean neural signals and dataset ***B*** contains subsequently-recorded, disrupted signals. The inputs ***x***_***A***_ and ***x***_***B***_ are unpaired individual clean and disrupted samples. Encoders embed their respective inputs into the latent space ***z***. Each decoder can operate on any point in the space ***z*** in order to reconstruct a signal of type ***A*** in the case of ***Dec***_***A***_ and of type ***B*** in the case of ***Dec***_***B***_. The latent representation of the sample ***x***_***B***_ that is decoded by ***Dec***_***A***_ results in the (fake) domain adapted signal ***x***_***BA***_. Each signal type has a discriminator that is trained to separate autoencoded signals from domain adaptations.

The neural network topologies we used for encoders and decoders were feed-forward, ReLU networks with dropout between layers. We did not apply activation functions or dropout in the final layer of the encoder in order to maintain a maximally expressive latent space, and we did not apply dropout before the final layer of the decoder in order to enable the greatest fidelity in reconstructed signals. Network topologies were fully-connected for spiking interfaces where there is no assumption of spatial correlation in signal space and convolutional networks were used for ECoG arrays since spatial correlation in those signal spaces may exist (particularly for mECoG where adjacent electrodes in the grid are separated by small distances). Discriminator networks were feed-forward, ReLU networks with a Sigmoid applied to the output layer, having either fully-connected or convolutional topology, again depending on the modality. Each discriminator operated on the signal space and produced a scalar output between 0 and 1, where Disc_A_(x_*_) > 0.5 is interpreted as ‘real’, or having come from the data collection, and Disc_A_(x_*_) <= 0.5 as ‘fake’, or having come through the recovery pathway Dec_A_(Enc_B_(x_B_)).

The architecture in [24] employed weight sharing in later encoder layers as well as in early decoder layers. We experimented with weight sharing in the encoder and observed no benefit.

### Training Objectives Rationale

Here we introduce the three categories of training objectives and their corresponding network weight updates in the order in which they are performed. For all weight updates we use PyTorch’s AdamW optimizer.

The significant difference between ANNI and the UNIT algorithm is evident here: UNIT uses statistical divergences in the latent space for both the autoencoding training objective (standard VAE) and in the cycle consistency training objectives, and ANNI does not. The reason for the difference is that, in our multi-class context, we have found that it is essential that each class is encoded into its own distinct region in the latent space in order to ensure that disrupted signals recover into the class that they correspond to. With ground-truth knowledge of signal class one could adopt statistical divergences conditioned on class in order to achieve the benefits of the VAE while also separating classes in the latent space (C-VAE literature), but this required class information is not available in our application.

### Autoencoder updates

The mapping x_AA_ ≔ Dec_A_(Enc_A_(x_A_)) is interpreted as a reconstructing autoencoder (as is Dec_B_(Enc_B_(x_B_))).

Loss function(s): We apply the signal reconstruction losses

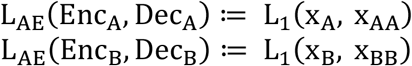

Weight updates: We update the weights in Enc_A_ and Dec_A_ based on the reconstruction loss L_AE_(Enc_A_, Dec_A_), and similarly we update the weights in Enc_B_ and Dec_B_ using the reconstruction loss L_AE_(Enc_B_, Dec_B_).

The purpose of these autoencoder training steps is to independently establish expressive feature extractors for non-disrupted and disrupted signals, as well as to establish capable decoders to independently reconstruct signals from the latent space.

### Discriminator and Generator updates

The discriminator and generator are trained in alternating steps. The first step updates each discriminator to learn the difference between incoming signals and recovered or artificially disrupted signals.

Loss function(s): We apply Binary Cross-Entropy as the training objective

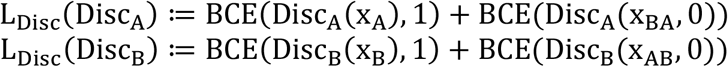

Weight updates: We update only the weights in the discriminators Disc_A_ and Disc_B_ at this stage. The second step updates the generators, which form the recovery pathway and the artificial disruption pathway. The recovery pathway is defined as x_BA_ ≔ Dec_A_(Enc_B_(x_B_)) and the artificial disruption pathway is x_AB_ ≔ Dec_B_(Enc_A_(x_A_)). We consider these to be ‘generative’ processes since the resulting signal is only statistically related to signals collected from the BCI, and has no counterpart in either dataset to match to.

Loss function(s): The adversarial loss for generators captures the extent to which the generator is fooling the discriminator.

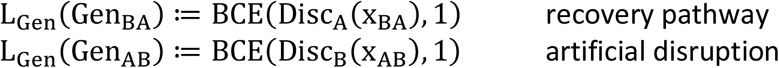

Weight updates: We update only the weights in the generator pathways at this step. The loss L_Gen_(Gen_BA_) is used to update the networks E_B_ and D_A_, while the loss L_Gen_(Gen_BA_) is used to update the networks E_A_ and D_B_.

The purpose of interpreting the recovery (and artificial disruption) pathway as generator, and guiding that generator using a discriminator, is to obtain a non-disrupted representation of each incoming neural recording that is statistically similar to the non-disrupted dataset. This does not solve the class matching problem, which we address that with Cycle Consistency and an ad-hoc drift penalty. This stage also includes an ad-hoc penalty on the dynamic range of the generator’s output. We have found, as is typical in many classification problems, that our discriminators are very tolerant of variance in the dynamic range of their inputs. Because we are trying to satisfy an arbitrary neural decoder, we aim to reconstruct disrupted signals as faithfully as possible. To this end we introduce:

Loss function(s): We apply the following signal-space penalties:

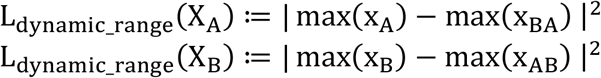

Weight updates: We update weights in encoders Enc_A_, Enc_B_ and decoders Dec_A_, Dec_B_ at this stage.

### Cycle Consistency updates

Cycle consistency describes the ability to accurately reconstruct a single incoming signal by first transferring it across one domain, and then transferring the result across the other domain. In symbols, starting with a single disrupted signal and finishing with what should be a very similar signal, this appears as

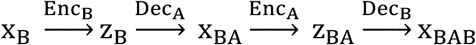

As pointed out by [24], two cycle consistency measures can be made from a single input x_B_ – one in the latent space: d(z_B_, z_BA_), and one in the signal space: d(x_B_, x_BAB_), where d is some distance metric. (Note: UNIT does not compare z_B_ to z_BA_ directly, rather they use the KL-divergence between a batch of latent vectors {z_B_} and the standard normal distribution, as well as the batch of latent vectors {z_BA_} and the standard normal distribution.)

Loss function(s): We apply loss functions in the latent space as well as in the signal space:

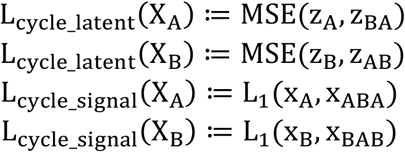

Weight updates: We update only the weights in the encoders Enc_A_ and Enc_B_ at this stage.

The importance of cycle consistency in our context is the opportunity it provides to align the latent representation of non-disrupted signals with disrupted signals from the same class. Cycle consistency itself does not guarantee class alignment alone, as seen by the following scenario that we encounter in the training and evaluation of ANNI. Below we introduce the recovery drift penalty L_drift_ to mitigate the undesirable class-swapping problem.

> **Example (class swapping):** Suppose x_B_ is from signal class 1, and consider the signal x_BAB_ that is the result of the cycle mapping of x_B_ that is drawn above. The training objective L_cycle_signal_(X_B_) is a signal-space comparison between x_B_ and x_BAB_, which implies that the optimized network will produce an x_BAB_ signal most similar to the class of its starting signal, which in this case is class 1.
>
> A problem can and does occur for the intermediate state x_BA_ of this cycle mapping – it may happen that either Enc_B_: x_B_ ↦ z_B_ encoded into to the wrong region in latent space, or that Dec_A_: z_B_ ↦ x_BA_ decoded into the wrong the class. In these scenarios, the training objective can still be fully optimized by guiding the second half the cycle mapping 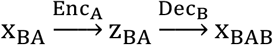 to undo the class switch.
>
> We call this class swapping and we have observed near-perfect signal recovery from very severe disruptions where the class identities of the recovered signals were a permutation of the class identities of the disrupted signals. This problem manifests visually as a permutation of the columns of a confusion matrix computed from the neural decoder’s classification of recovered signals x_B_.

To directly address the class swapping problem, we penalize deviation in signal space between the input signal and the recovered (symmetrically the artificially disrupted) signal. The intuition here is that disrupted signals are closer to non-disrupted signals of the same class than they are to non-disrupted signals of any other class.

Loss function(s): The following signal-space penalties are applied during training:

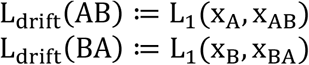

Weight updates: We update weights in encoders Enc_A_, Enc_B_ and decoders Dec_A_, Dec_B_ at this stage.

### Training

The system is trained using two datasets: X_A_ consisting of unlabeled non-disrupted neural recordings from many classes (up to 20 in our simulations); and X_B_ consisting of unlabeled disrupted neural recordings from the same signal classes as X_A_. No pairing of samples is possible in this setting. Batches of sample pairs ({x_A_}, {x_B_}) are randomly drawn and used to feed the network mappings described in Section 0 over anywhere between 100 and 1000 epochs.

We consider our training to be complete when the decoder entropy on the disrupted set of signals is at a maximum (taken over some reasonable, pre-defined number of epochs). Decoder class entropy is computed at the end of each training epoch by collecting the neural decoder prediction of each recovered neural signal x_BA_, recovered from the set X_B_. Figure 4 shows a typical history of decoder entropy along with the decoder accuracy during the epoch training of a single model. As decoder accuracy on disrupted signals is not accessible at all during the training and operation of ANNI, decoder entropy is a valuable surrogate.

### Application of a trained model

Signal recovery is the raison d’être of the ANNI system. The simplest application of a trained ANNI model is to pass each incoming, presumably disrupted, neural recording x_B_ corresponding to a class that is decodable by the neural decoder into the recovery pathway:

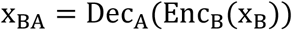

This signal is then available to be passed directly to the neural decoder as the next step in actualizing the intended benefit of the BCI.

## Results

### MEA dataset decoder recovery

MEA recordings can be disrupted either by losing neuron channels completely (“dropped”) or by changes in the neuron channels (“swapped”). Increasing the number of dropped or swapped channels increases the chances for decoder error and increases the challenge of data recovery. Thus, ANNI’s ability to recover decoder accuracy was tested using datasets with increasing numbers of dropped or swapped channels of PSTH data. For each disrupted dataset, 5-10 different ANNI recovery models were trained and one model was identified using the entropy selection criteria (see Methods). Recovery was tested at the same level of disruption the model was trained, but selected models are also placed into competition with newly trained models at the subsequent disruption step.

ANNI recovered MEA datasets had significantly higher decoder accuracy than unrecovered drop-out datasets. Decoder accuracy on unrecovered disrupted data decreased with the number of MEA channels dropped (red points and line). In contrast, decoder accuracy varied widely for ANNI recovered data (grey points), even for the same models instantiated with different random seeds. However, recovered datasets from entropy selected models (blue circles) outperformed the majority of models and outperformed the unrecovered data in almost all datasets. Indeed, ANNI recovered datasets, maintained decoder accuracy >90% in all datasets with as much as 13 of 32 channels dropped. In the majority of cases newly trained models outperformed previously selected models but in five cases the previously selected model was selected again (blue lines). Similar results were observed when the neurons’ tuning curves and order of channels dropped was changed (data not shown).

ANNI was also successful in recovering data when MEA channels were swapped instead of dropped (see Figure 6). Similar to decoder performance when MEA channels were dropped, decoder accuracy on unrecovered data decreased with the number of channels swapped (red points and line) and varied widely for recovered datasets (grey points). However, recovered datasets using entropy selected models (blue circles) outperformed the unrecovered data in all tested disrupted datasets. ANNI recovered datasets maintained decoder accuracy >90% in 21 of 31 datasets. Notably, ANNI recovered a dataset to >90% accuracy with 30 of 32 channels swapped and an unrecovered decoder accuracy near 20%. Similar results were observed when the neurons’ tuning curves, the order channels swapped and the direction of swap (e.g. increase vs. decrease in tuning width) was changed. These results illustrate ANNIs potential to recover neural data from a range of disruption types and datasets.

**Figure 6.**
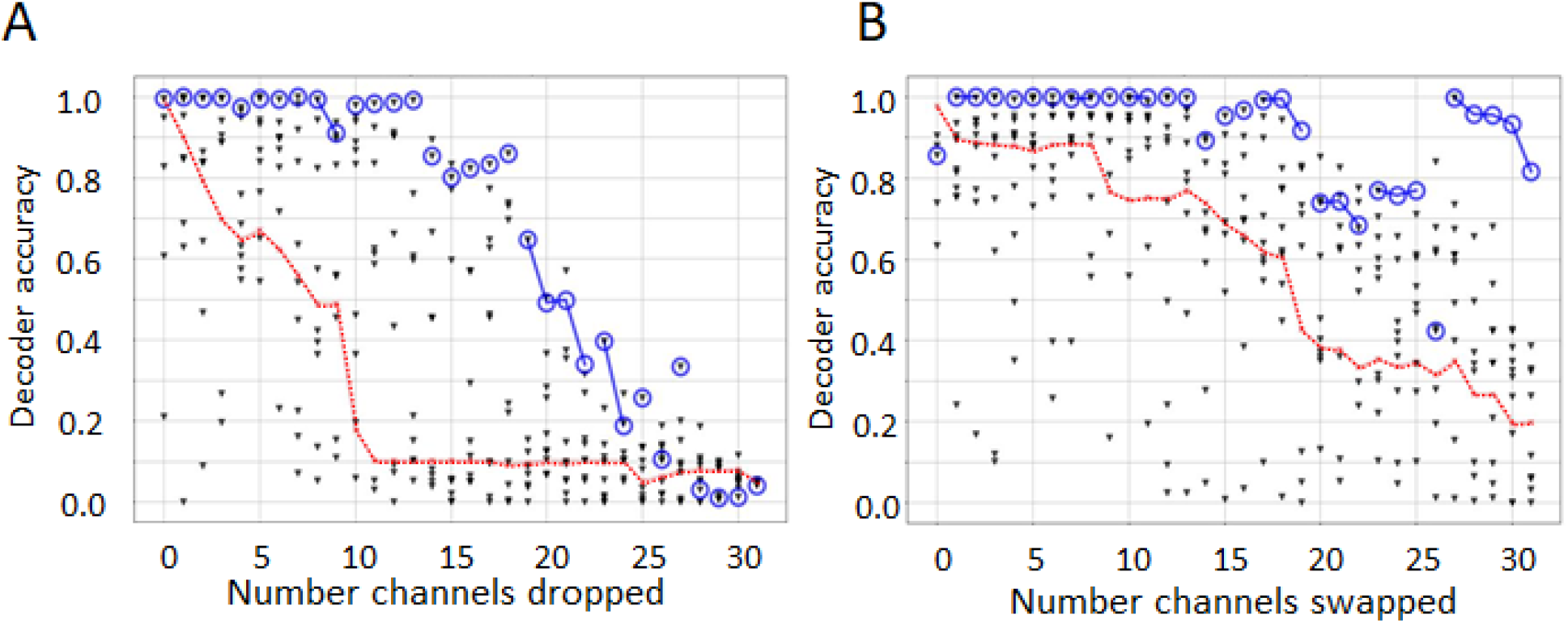
The decoder accuracy of recovered datasets from selected ANNI models substantially outperforms most unrecovered datasets and non-selected ANNI models. **(A)** Decoder accuracy declines with the number of dropped channels in the unrecovered PSTH datasets (red lines). The decoder accuracy of datasets recovered with ANNI models (grey points) is highly variable but entropy-selected models (blue circles) generate some of the highest decoder accuracies of all possible models. Blue lines are shown when a previously selected model was maintained as the highest entropy model on the subsequent disruption step. **(B)** Similar results are observed with swapped disruption.

### ECoG dataset rescue

We have tested the ability to recover from ECoG disruption simulated at 10-day increments from 10-360 days. Figure 7 shows the performance of ANNI across the simulated lifetime of a chronic implant. The red dashed line indicates the performance of the decoder on the raw degraded data and shows how quickly the accuracy degrades in the first 100 days. Consequently, this is the reason why many investigators using chronically implanted ECoG do not train neural decoders until after the first 70-100 days when signals have stabilized. From this point, variability is observed in the accuracy of the decoder reflective of the random changes in signal strength over the remainder of the days after implant. The gray dots at each assessed time point indicate multiple ANNI models which were trained/tested for their accuracy but not selected by our current model selection criteria, decoder entropy (see Methods), as the model that was used. The blue circles indicate decoder performance on the selected ANNI models (automated entropy criteria). ANNI nearly completely restores the performance of the decoder at every time point after implant. At most time points our model selection criterion selects the best model and, in all cases, it selects models in the top 3% of trained models. ANNI is incredibly successful at signal rescue in these interfaces. The success likely stems from the fact that the information contained in the signals is relatively stable, although the SNR is highly variable across samples. These shifts in signal amplitude, even when experienced unevenly, are possible to translate back with relative ease by the algorithm.

**Figure 7.**
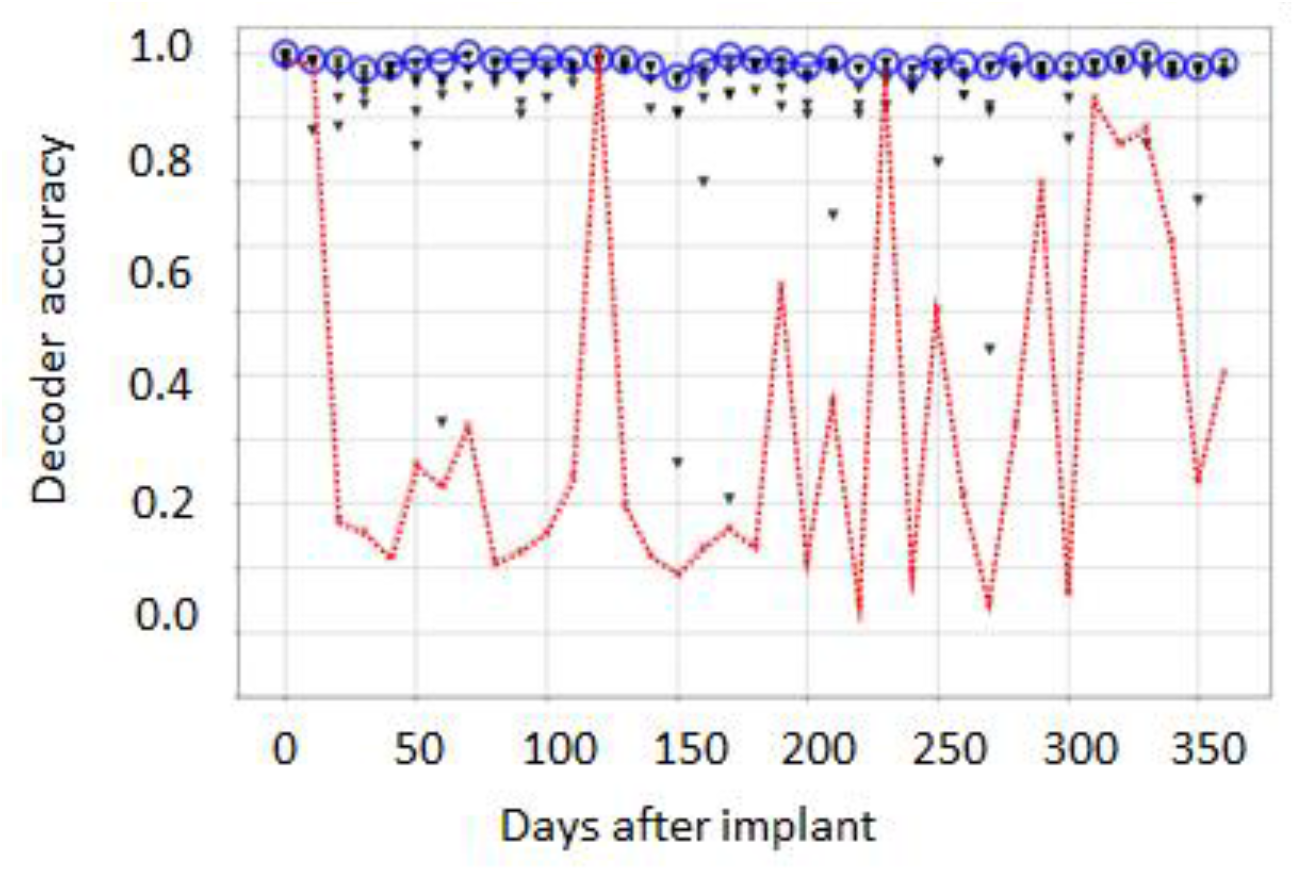
ANNI consistently restores signals to improve decoder accuracy in the face of catastrophic and serial disruption. The red dotted line shows the performance of the decoder on the raw degraded data at each step (decoder trained at day 0). The blue markers/line show the performance of data pushed through selected recovery models. Model selection done using previously described criterion of entropy. Gray dots indicate performance of trained but unselected recovery models.

## Discussion

We have developed a GAN based network, ANNI, to maintain decoder stability during signal disruption. While adapted from a UNIT architecture, ANNI was modified in several important ways to remap neural signals from the disrupted to clean domains without loss of “class” information that would cause errors in the neural decoder. Most prominently, ANNI does not utilize a variational autoencoder (VAE) but uses a cycle-consistency latent space training objective that encourages data from different domains but the same “class” to map consistently in the latent space. We note that the maintenance of “class” identity across domain remapping is similar to the maintenance of “content” across “style” domains performed by MUNIT, itself a derivation of the UNIT architecture [30]. While similar in function, ANNI is less complex in structure (e.g. ANNI uses a single shared latent space) and we are encouraged by ANNI’s performance remapping the simulated BCI data.

ANNI demonstrated the ability to recover decoder performance from two different simulated types of BCI with distinct types of signal disruption. ANNI was able to maintain decoder performance near 100% accuracy when 13/32 recording channels were dropped or 27/32 channels were swapped. Without intervention from ANNI, dropping 13/32 or swapping 27/32 caused the decoder performance to fall near 10% and 30% respectively. With minor changes to the network, ANNI was adapted to remap simulated ECOG data instead of MEA data. In the case of simulated ECOG signals, ANNI-rescued data maintained decoder performance near 100% accuracy, while decoder performance on control disrupted signals fell below 20%. These results demonstrate ANNI’s potential to maintain decoder accuracy in distinct BCI types during diverse signal disruptions. Below, we put these results into context of other techniques designed to maintain BCI decoder accuracy and suggest future directions to utilize ANNI.

ANNI and other recent techniques to stabilize BCI decoders leverage the inherent stability of low-dimensional BCI data. BCI data is often high-dimensional, having many recorded channels and time bins. However, while neural measurements are often high-dimensional, the information can be well represented in a low-dimensional space [32]. ANNI effectively learns low-dimensional representation of both the clean and disrupted neural data by training a set of autoencoders with a limited latent space to reproduce the data. Further, the successful remapping across data domains relies on the similarity between disrupted and clean latent representations. Thus, ANNI relies on the fact that while the high dimensional neural measurement may be significantly changed by the disruption, its lower dimensional representation often remains stable. Other work has leveraged lower dimensional representations of neural data to increase BCI stability [33-35]. Most notably, Degenhart and colleagues used low-dimensional manifold representation of BCI data to maintain stability in the face of BCI signal disruptions [34]. Similar to ANNI, manifold-based stabilization was successful in promoting decoder stability via a low-dimensional representation but differs significantly from ANNI in important ways. First, unlike ANNI, the manifold alignment requires methods for identification of stable electrodes and linear manifolds. On the other hand, ANNI requires decisions on network options, as exemplified by distinct networks for MEA and ECOG datasets, and post-training selection criteria. While we found success using the described networks and entropy selection criteria, we do not rule out possible improvements with other choices. Additionally, the decoder used by Degenhart and colleagues was trained to latent space representation, thus coupling the BCI decoder to the manifold-based stabilization algorithm. In contrast, ANNI recovers signals to the original BCI input space, thus allowing ANNI to be used with any decoder trained on the original BCI data with limited or no knowledge of the decoder itself. Thus, while ANNI leverages the stability of low-dimensional representation of BCI signal, similar to other stability algorithms, there are important differences in how these low-dimensional representations are found and utilized.

Beyond algorithms to improve fixed decoder stability, recent work has developed techniques to minimize the time and effort necessary to retrain a BCI decoder [36-39]. These techniques show significant improvements over previous training regimes but may still require significant user engagement [34]. However, because ANNI does not alter the signal structure from the BCI (e.g. number of channels or time bins) it can be incorporated with other techniques. Thus, techniques to speed retraining of BCIs could use ANNI to further minimize the error and user time involved in retraining the decoder or BCI user.

While the experiments described above demonstrate ANNI’s potential to successfully maintain BCI decoder accuracy, there are several important experiments needed to confirm ANNI’s utility. First, ANNI must be tested with non-simulated longitudinal BCI data and a fixed decoder. While the simulated BCI data used in this paper was designed to mimic key aspects of MEA and ECOG measurements, neural responses are often highly complex (e.g. firing rate and correlation structure temporal dynamics) and differ greatly across brain region; Such diversity and complexity was not necessarily captured by our simulated data. Second, ANNI’s training limitations must be better assessed. The experiments described above trained ANNI with 5000 samples of clean and disrupted data. Depending on the rate of disruption and data collection, such large sample numbers may not always be practical. While pilot experiments on BCI data from rats suggest ANNI can perform successfully outside of our simulated datasets, these experiments need to be more rigorously tested.

While more work needs to be done to confirm ANNI’s utility we are encouraged by the results reported in this paper. BCI technology and decoder performance has greatly improved since early experiments but its practical utility remains substantially limited in large part due to the instability of the BCI and decoder. ANNI provides a potential path to prolong BCI functionality and enable its growing role in therapeutic intervention.

## Acknowledgments

The authors would like to acknowledge Dr. Robert Hampson and Brent Roeder at the Wake Forest Baptist Medical Center for their help in understanding signal disruption on MEAs.

## Footnotes

1 Up to four models with relatively large datacollects of 5,000 samples can train simultaneously on a single GPU (NVIDIA GTX 1080 or better) within an hour or two. Signal array size and network architecture are the most important factors to training time.

